# Bioluminescent Genetically Encoded Glutamate Indicator for Molecular Imaging of Neuronal Activity

**DOI:** 10.1101/2021.06.16.448690

**Authors:** E. D. Petersen, A.P. Lapan, A.J. Fillion, E. L. Crespo, G. G. Lambert, A. Torreblanca Zanca, R. Orcutt, U. Hochgeschwender, N. C. Shaner, A. A. Gilad

## Abstract

Genetically encoded optical sensors and advancements in microscopy instrumentation and techniques have revolutionized the scientific toolbox available for probing complex biological processes such as release of specific neurotransmitters. Most genetically encoded optical sensors currently used are based on fluorescence and have been highly successful tools for single-cell imaging in superficial brain regions. However, there remains a need to develop new tools for reporting neuronal activity *in vivo* within deeper structures without the need for hardware such as lenses or fibers to be implanted within the brain. Our approach to this problem is to replace the fluorescent elements of the existing biosensors with bioluminescent elements. This eliminates the need of external light sources to illuminate the sensor and overcomes several drawbacks of fluorescence imaging such as limited light penetration depth, excitation scattering, and tissue heating that are all associated with the external light needed for fluorescence imaging. Here we report the development of the first genetically encoded neurotransmitter indicators based on bioluminescent light emission. These probes exhibit robust changes in light output in response to extracellular presentation of the excitatory neurotransmitter glutamate. We expect this new approach to neurotransmitter indicator design to enable the engineering of specific bioluminescent probes for multiple additional neurotransmitters in the future, ultimately allowing neuroscientists to monitor activity associated with a specific neurotransmitter as it relates to behavior in a variety of neuronal and psychiatric disorders, among many other applications.

## Introduction

Optical biosensors have proven incredibly useful to researchers in a wide variety of fields for the study of neuronal and cellular activity^1,2^. These probes generate changes in light emission intensity and/or wavelength in response to physiological events such as calcium influx, membrane voltage changes, or the presence of a ligand such as a neurotransmitter^3–7^. Currently, nearly all available biosensors rely on fluorescence, leaving much room for improvement and further development of new approaches to better report changes in these cellular dynamics. One major improvement to such indicators would be to eliminate the need for an excitation light source, which is necessary to excite fluorescent reporters.

The illumination source used for fluorescent imaging is a major limiting factor for the depth at which cells can be imaged through tissue. This is due to the scattering of light traveling into the tissue, as well as heating as the incident photons are absorbed by endogenous molecules in the tissue. Imaging with bioluminescent probes eliminates the need for excitation light, circumventing these issues and enabling researchers to image deeper structures of the brain^8^. Furthermore, autofluorescence produced by the illumination sources used for fluorescent imaging is not present when using bioluminescence, allowing for enhanced signal detection in deeper structures since the signal to noise ratio can be higher. These fundamental limitations to fluorescence imaging and corresponding benefits of bioluminescence imaging dovetail with an urgent need in the field of neuroscience: tools that allow recording and modulation of entire neuronal populations that are both non-invasive and don’t require implanted hardware. (Fig. 1).

**Figure 1.**
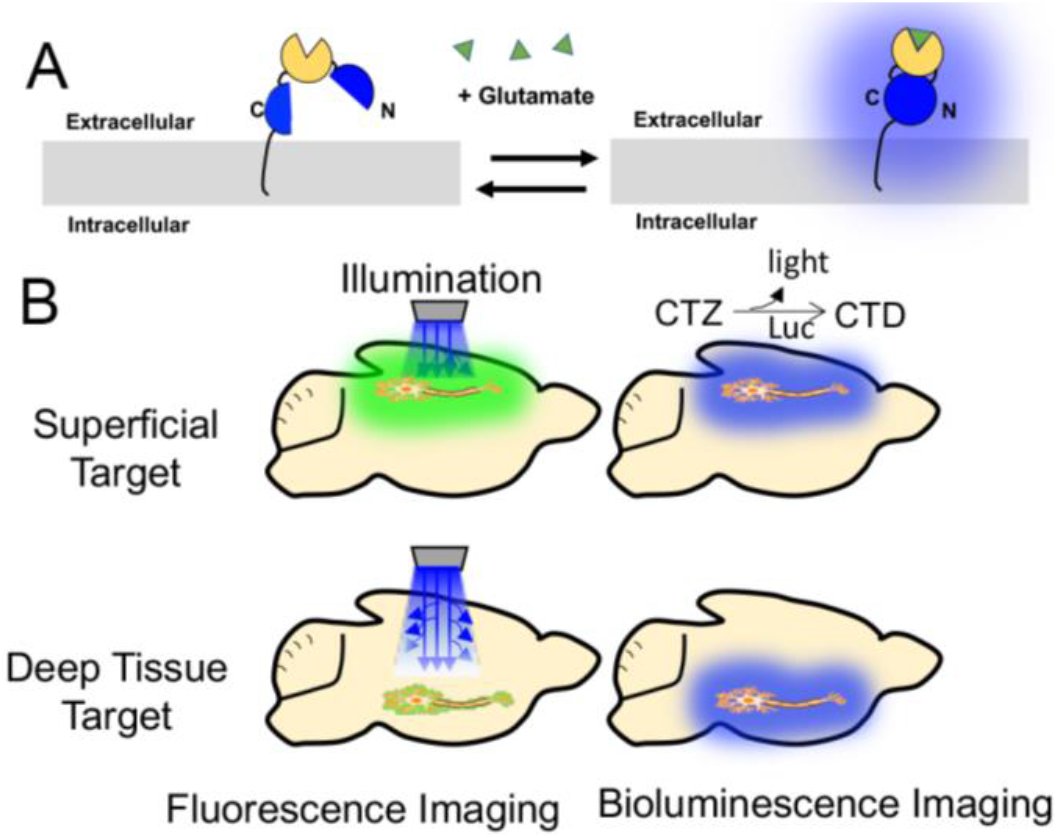
A. Schematic of the BioLuminescent Indicator of the Neurotransmitter Glutamate (BLING) with a split luciferase (blue) and glutamate sensing domain (tan) displayed on the cell surface. B. Comparison of bioluminescence vs fluorescence imaging for superficial and deep tissue targets in the rodent brain. Fluorescence requires an excitation light which scatters as it interacts with the tissue, decreasing the ability to efficiently excite indicators as depth is increased. Bioluminescence is produced within the tissue through an enzymatic reaction of a luciferase (Luc) with its substrate (e.g. coelenterazine, CTZ or furimazine) to produce light and an oxidized by-product (e.g. coelenteramide, CTD or furimamide), which allows for increased imaging depth as an excitation light source is not the limiting factor.

Bioluminescence is produced by an enzyme (luciferase) that catalyzes oxidation of its specific substrate called a luciferin, resulting in the emission of light. Different forms of biological light production exist in multiple domains in nature, including beetles, worms, bacteria, and the majority of marine organisms^9,10^. Bioluminescence has been used for a variety of imaging applications, such as the quantification of gene expression over time and for imaging calcium dynamics to represent neuronal function or to track other calcium events within cells.^11–13^ Although bioluminescence has been used in research for decades, more recently scientists are rapidly improving luciferases and synthetic luciferins. For example, a 1000-fold increase in luminescence was recently achieved by evolving firefly luciferase to use a synthetic substrate to create AkaLuc which produces near infrared bioluminescence.^14,15^

Before genetically-encoded indicators were available, the standard method to analyze specific neurotransmitters *in vivo* was to collect cerebrospinal fluid from within the brain via microdialysis or cyclic voltammetry, both of which offer poor time resolution (multiple minutes) and have decreasing performance in longitudinal studies. In the past several years, fluorescent genetically encoded neurotransmitter indicators such as iGluSnFr, Dlight and GRAB-DA^7,16,17^ have enabled detection of neurotransmitters on a faster time scale. Unfortunately, fiber photometry, the implantation of an optical fiber into the brain, is required to measure the output of these fluorescent probes in deep regions of the brain. To address these shortcomings of current methods, we are developing a series of bioluminescent Genetically Encoded Neurotransmitter Indicators (bGENIs) to provide neuroscientists with a set of tools that can be used in lieu of physical collection and fluorescence detection approaches. We have made significant progress in developing a BioLuminescent Indicator of the Neurotransmitter Glutamate (BLING). Glutamatergic neurotransmission is directly implicated in behavior, movement, mental health, pain perception and addiction, making it an attractive target for the development of a bGENI. BLING will also have a variety of drug discovery applications for use in high throughput screening. In the future, the design and engineering approach we describe here can be expanded to encompass all neurotransmitters as well as other small-molecule analytes. Moreover, the BLING sensor and our engineering approach can be further adapted to create novel therapeutic agents for a variety of neurological disorders by acting on light-sensitive proteins (optogenetic actuators).

## Results

To expand the neuroscientists’ toolbox of genetically encoded indicators, we have engineered the first reported bioluminescent genetically encoded neurotransmitter indicators using a multistep screening approach. In the first step, sensors were rationally designed using a variety of split luciferase variants to fuse the luciferase halves to a sensing protein. Then, we devised an automated workflow to screen for improved variants in mammalian cells. We initially created and tested 3 versions of BLING constructs based on the truncated periplasmic glutamate binding protein (Glt1) from the FRET-based glutamate sensor SuperGluSnFr^18^ flanked by short flexible linkers with a split luciferase “half” on each terminal. Initially, we chose various marine luciferases that have been engineered into ultra-bright variants, do not require a cofactor to produce light, and have previously validated split sites. We created BLING 0.1 consisting of the *Gaussia* luciferase (GLuc) variant M43L, M110L, referred to as “Slow Burn GLuc,” which is brighter than native GLuc and has glow kinetics instead of flash kinetics.^19,20^ We used the split site 105-106 previously used to successfully engineer calcium indicators^21^, including the 17 AA native secretion signal from GLuc for surface display with a PDGFRβ membrane anchor. BLING 0.2 was created similarly using NanoLuc split at 66-67 which has been used to successfully engineer calcium indicators^22^, with an Igk leader sequence for cell surface display. BLING 0.3 was created with NanoLuc large and small bits split at 159-160, which has also previously been used to generate calcium indicators^23^. All initial BLING variants produced bioluminescence with native CTZ for Gluc or h-CTZ for Nluc with BLING 0.2 having the largest response to glutamate and being the brightest sensor. (Fig. 2A,C)

**Figure 2.**
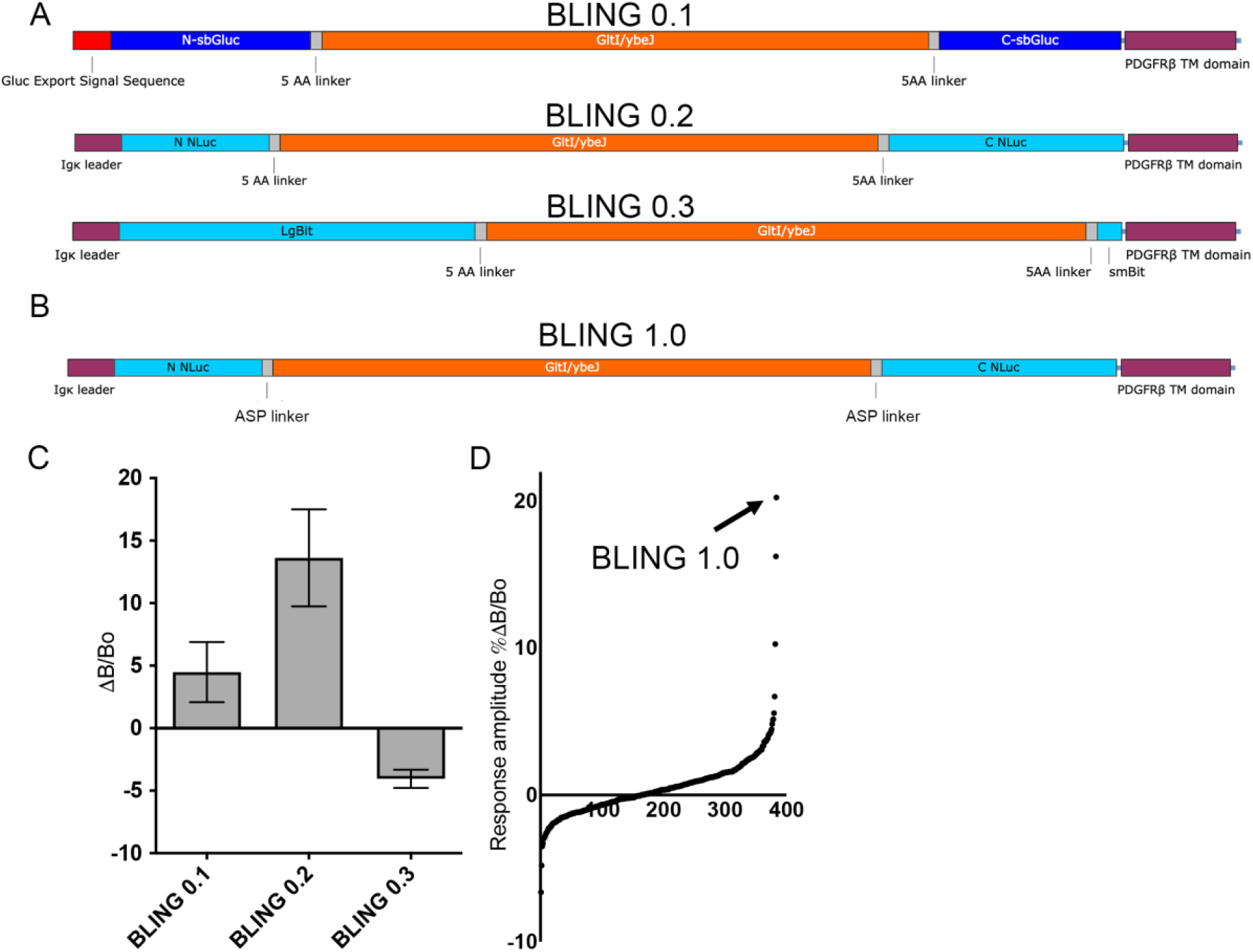
A. Protein maps of the three initial BLING designs that were tested. From top to bottom: slow burn *Gaussia* luciferase split at 105-106 including the native *Gaussia* secretion peptide on the N terminal instead of the IGK leader secretion sequence; Nanoluc split at 66-67, Nanoluc split at 159-160 all with Glt1 glutamate binding protein and PDGFRβ transmembrane domain to anchor the sensor to the extracellular side of the membrane. B. Protein map of BLING 1.0 with variable linkers. C. Results from the initial BLING constructs in response to 1 mM glutamate. n=4. D. Responses from the BLING linker library of ~400 variants of BLING 0.2 with variable linkers tested.

Following the testing of our initial 3 variants we aimed to improve the top performing BLING design through linker optimization. BLING 0.2 was selected for further engineering and the linkers were replaced with 3 amino acid variable linkers that code for A, S or P at each position (Fig. 2B, Sup. File 1). This combination of linker variants was used because they have been shown to produce diverse functionalities due to varying levels of rigidity/flexibility while significantly limiting the number of clones that need to be screened^23^. From this library, the top performing variant was selected for further characterization (Fig. 2D). As a result of linker optimization, we were able to generate a BLING variant (BLING 1.0, Addgene plasmid: 171647) that has a robust response to glutamate addition when expressed in mammalian cells (Fig. 3B). BLING 1.0, derived from BLING 0.2, consistently outperforms the parental construct by 2-fold in terms of response to glutamate while maintaining its brightness using multiple plate reader modalities. However, the response amplitude can vary significantly depending on plate reading modality (Figs. 3C and S1). We also found both BLING variants to outperform iGLuSnFr in terms of magnitude of percent change in response to 1 mM glutamate and GCaMP6m’s response to 5 μM ionomycin when doing bulk measurements of the whole wells light emission using a plate reader (Fig. 3B). We next sought to determine the dose dependent response of BLING 1.0 when expressed in mammalian cells using a photon counting plate reader (Tecan Spark). The responses to background levels of glutamate were minimal, and we observed a 10.8% change in response to 10 μM with a threefold greater response to 100 μM with a 31.8% change in luminescence (Fig. 3C).

**Figure 3.**
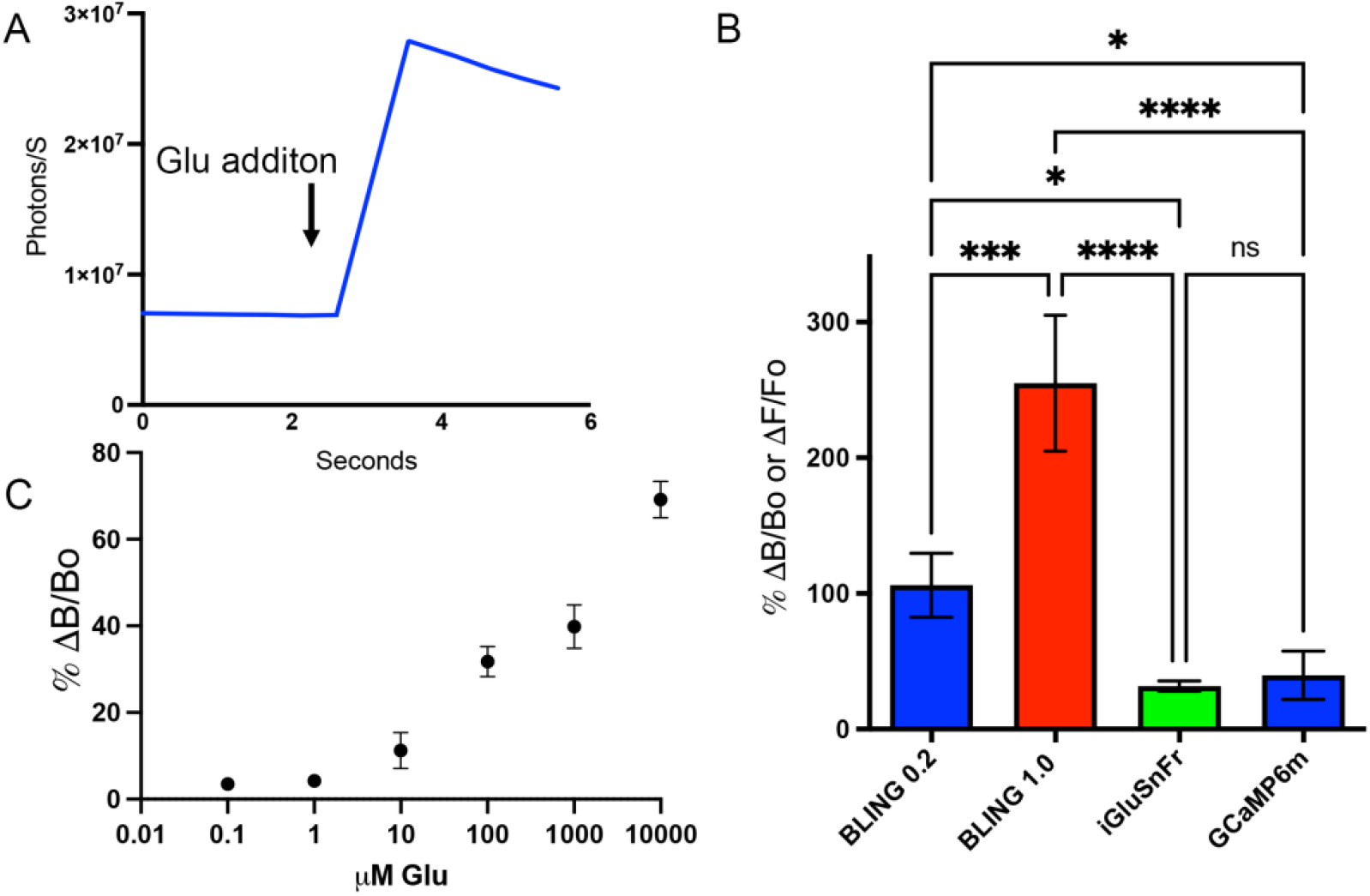
A. Example responses of BLING 1.0 to 1 mM glutamate addition recorded with a plate reader. B. Response of the parental bling (BLING 0.2) compared to optimized BLING (BLING 1.0) and the fluorescent glutamate indicator iGluSnFr to 1 mM glutamate addition and GCaMP6m to 5 μM ionomycin taken as bulk measurements of the cell cultures with plate readers, Tecan Spark for BLING and Biotek Cytation 5 for fluorescent readings. n=2 for BLING, 3 of iGluSnFr, 12 for GCaMP. C. Dose dependent response of BLING when using a plate reader. n=3. *=p<0.05, **=p<0.01, ***=p<0.001, ****=p<0.0001.

Since BLING 1.0 successfully reports changes in extracellular glutamate levels when recorded on the population level we next sought to determine if changes in extracellular glutamate can be observed at the single cell level. For this we used live cell bioluminescence microscopy using HEK cells expressing BLING and perfusing varying concentrations of glutamate into the cell imaging chamber. We found BLING 1.0 to report changes in extracellular glutamate with responses up to 310% at the single cell level to 1 mM glutamate, average response of 109.6% (Fig. 4A, C and supplementary movie). We also determined that this sensor reports glutamate in a dose dependent manner, responding to physiological levels of glutamate and slightly outperforming BLING 0.2 (Fig. 4B).

**Figure 4.**
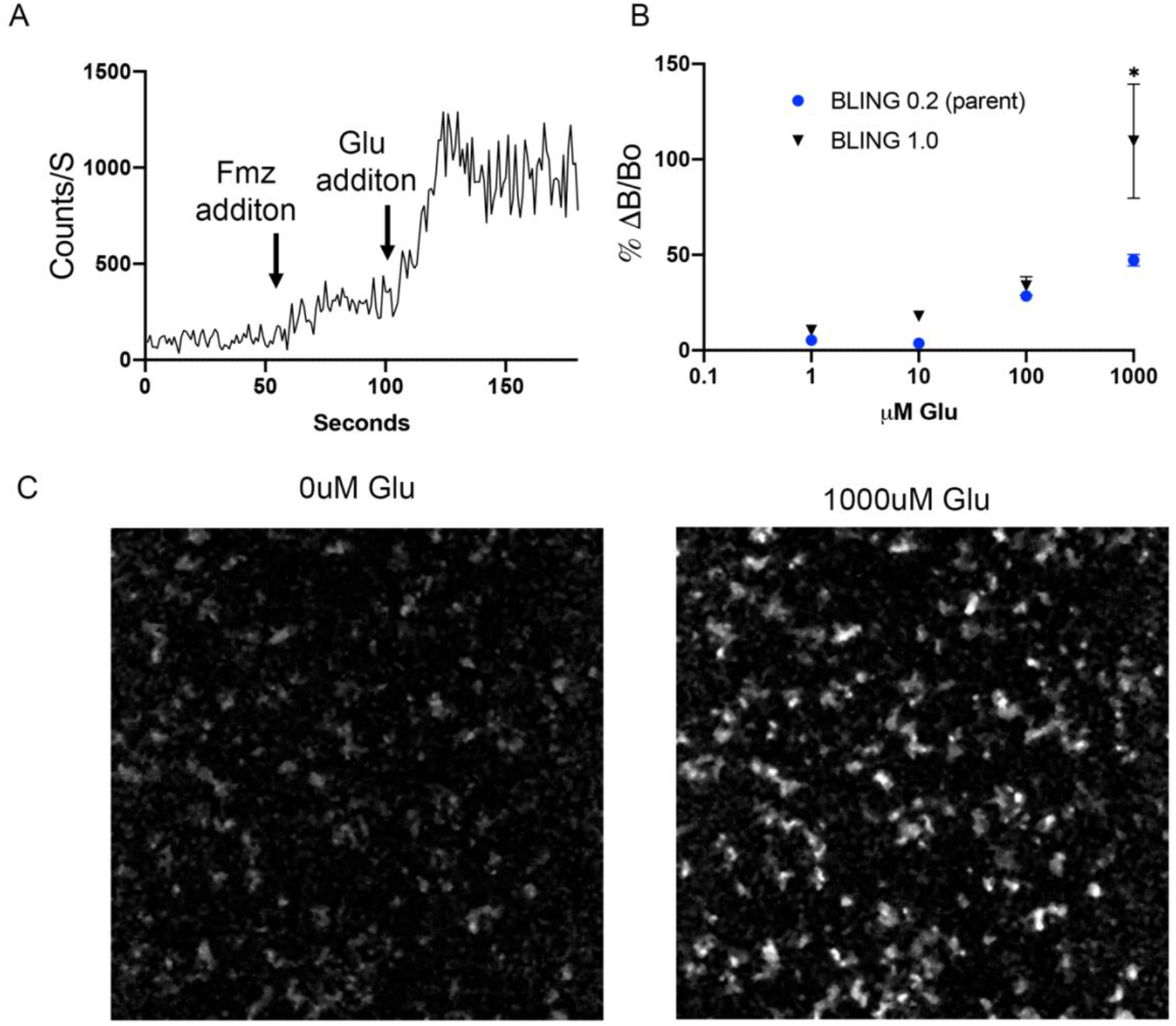
A. Example trace of a single cell ROI showing the perfusion of Furimazine starting at 60 sec and the infusion of 1 mM glutamate at 120 sec. B. Dose responses of BLING 0.2 and the improved variant BLING 1. N=4 for BLING 0.2, 8 for BLING 1.0 C. Image of background bioluminescence of BLING 1 and with 1 mM glutamate. *=p<0.05

We next sought to determine if BLING 1.0 can successfully report changes in glutamatergic activity *in vivo* by with an acute seizure model induced by local bicuculine injection. BLING 1.0 was expressed with AAV in the sensory cortex 2 mm below the surface of the skull (Fig. 5A). For imaging, h-CTZ was delivered intraperitoneally and rats were imaged with an IVIS Spectrum. In this preliminary proof of concept experiment we were able to detect increases in luminescence following seizure induction in four out of six rats (Fig. B,C) with increase in luminescence over 100%.

**Figure 5.**
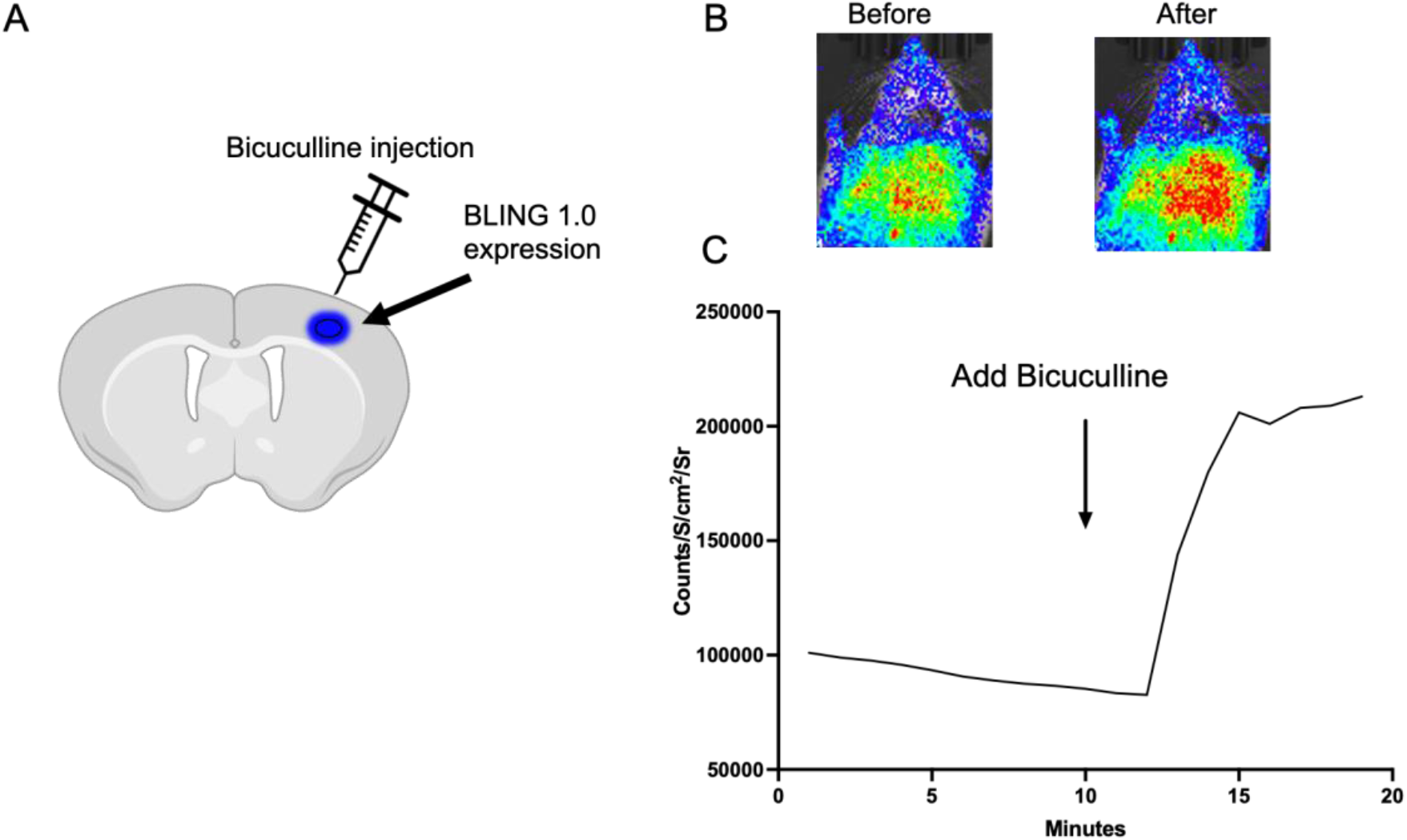
A. Schematic of the experimental design for expression of BLING 1.0 (blue circle) 2 mm deep within the sensory cortex, with bicuculline injected locally to induce an acute seizure. B. Example images of bioluminescence before and after seizure induction. C. Example trace of bioluminescent intensity in response to acute seizure induction.

In an effort to continue improving the BLING family of sensors we sought to create variants with lower levels of background luminescence and greater responses to glutamate. For this we created and tested BLING variants containing either flexible or rigid linkers in combination with three different variants of the small C terminal fragment, SmBiT with the large N terminal fragment that was evolved for improved protein stability previously described^24^ to create twelve new variants. This approach was based on our results from the randomized linker screening used to generate BLING 1.0 where we discovered that the top five variants generated all contained similar linker compositions with the first linker being composed of amino acids with flexible properties and the second linker containing primarily prolines with rigid properties. We also tested three C terminal peptide variants of NanoLuc in combination with the flexible or rigid linkers which have varied affinities to the N terminal of the luciferase ranging from ~1 nM to ~200 μM. We found that the variant described in ^24^ which has the highest affinity (peptide 86) resulted in four BLING variants that all responded to glutamate with 1.5-2 times the delta as BLING 1.0 with 1mM glutamate while having lower background light emission prior to glutamate addition with minimal effect from linker composision. We found BLINGs created with the other two peptides, the native NanoLuc peptide and peptide 114, the medium lowest affinity variants all had lower responses to glutamate presentation but were greatly affected by the linker composition (Fig. 6 and sup Fig. 1). BLINGs containing the native peptide with either flexible-flexible or rigid-flexible linkers compositions had 0.61 and 0.79 times the response of BLING 1.0 while variants containing peptide 114 with flexible-flexible or rigid-flexible linkers had 0.46 and 0.49 times the response and much lower background luminescence. Variants containing the native NanoLuc peptide or peptide 114 with rigid-rigid or rigid-flexible linker compositions all performed poorly. As can be seen in Fig. 6 and sup. Fig. 1, we created a new sub-family of BLINGs that is based on LgBiT ans SmBiT variants with lower luminescence background yet display greater increase in luminescence upon glutamate binding. Some of these BLINGs may prove more useful for imaging as they are still very bright and have greater responses while some variants may not have any use for imaging given that they are much dimmer but could be useful as optogenetic actuators since they may be able to avoid activation of light sensitive elements in their off state. Taken together, we have described here, a new family of synthetic biosensors that allow glutamate visualization both *in vitro* and *vivo* and have created a diverse subset of sensors that can be used as new starting points for further evolution and since these sensors produce their own light, they may have applications beyond neurotransmitter sensing.

**Figure 6.**
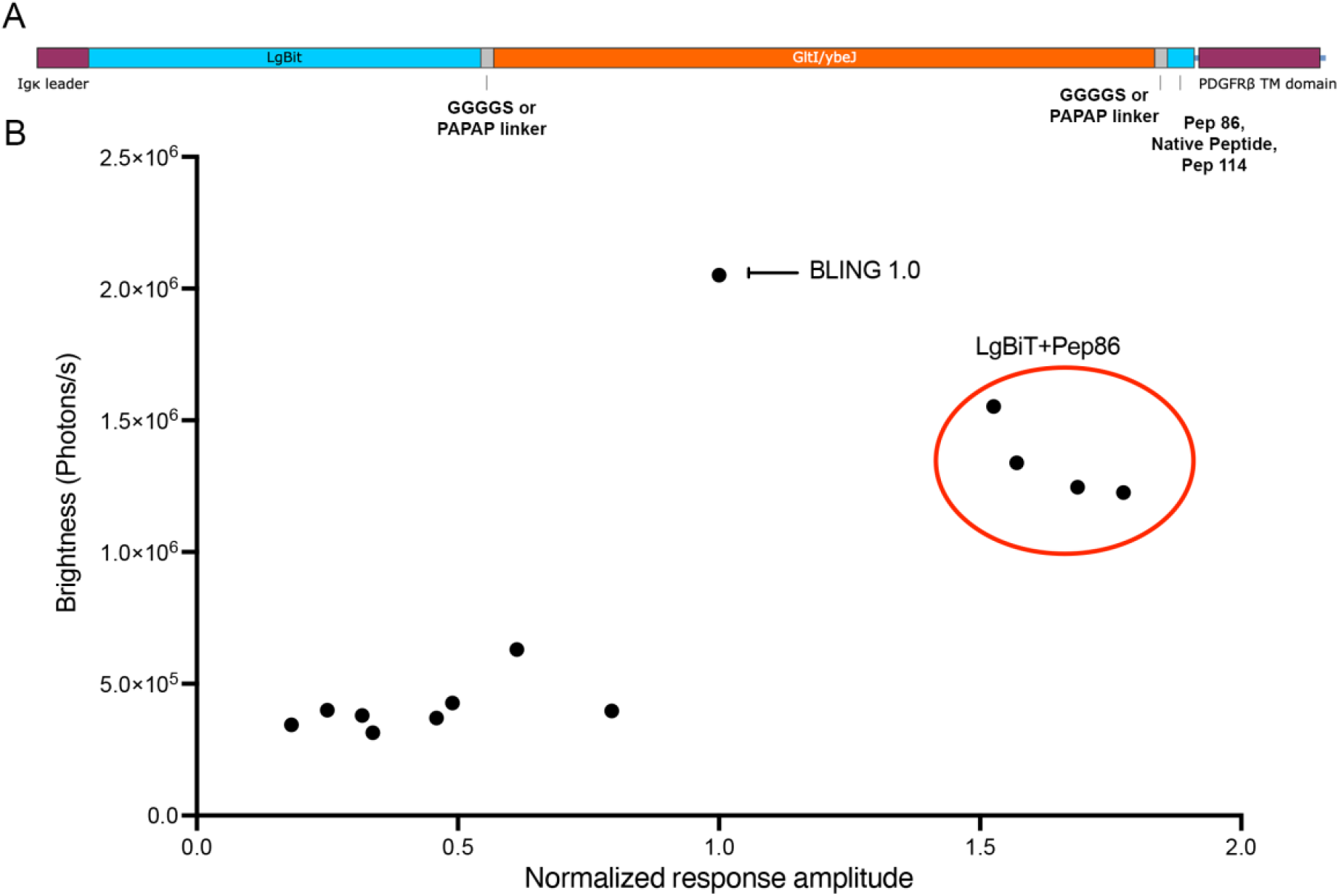
A. Protein map of the twelve new BLING variants that were tested that contain the 11s variant of NanoLuc called LgBiT for the N terminal half of the luciferase with either a flexible or rigid linker flanking the Glt1 glutamate binding protein and three C terminal variants with varying affinities to LgBiT Peptide 86, native peptide or pep 114 and PDGFRb transmembrane domain to anchor the sensor to the extracellular side of the membrane. B. Scatter plot of the diversity of new BLING constructs response to glutamate normalized to BLING 1.0 vs baseline brightness.

## Discussion

Here we present the first genetically encoded bioluminescent neurotransmitter indicator, which reports changes in extracellular glutamate via changes in luminescence intensity. This was achieved by building on previous work done to engineer various fluorescent glutamate indicators based on the same periplasmic glutamate binding protein (Glt1), SuperGluSnFr and iGluSnFr^7,18^. In this study, we replaced the two fluorescent proteins from SuperGluSnFr with N and C terminal fragments of various marine luciferases. These indicators work well to report changes in extracellular glutamate when expressed in cultured cells. This new tool opens the possibility for high throughput drug screening in cells as our bioluminescent indicator is ultra-sensitive when used in a plate reader compared to fluorescent indicators which do not perform as well when recorded at the level of whole cell cultures i.e. in bulk measurements. In addition to neuronal recording, we expect BLING to be extremely useful in situations where fluorescent indicators can’t be used such as when screening drugs or compounds that are optically active, for example where the drug itself is fluorescent.

The major advantage that bioluminescent indicators such as BLING present over currently existing fluorescent indicators is that they produce their own light, not requiring an excitation light source. Excitation light can become problematic if researchers don’t carefully consider the intensity of illumination, such as increasing illumination power when attempting to image deep brain areas. Overpowered excitation light can alter neuronal activity, heat tissue, lead to activation of astrocytes and microglia, cause scaring and cell death^25–30^. An additional constraint of fluorescence imaging is that many of the best fluorescent indicators currently used for optically recording neuronal activity suffer from significant photobleaching, often severely limiting the amount of continuous recording times to less than an hour in most cases.^31–34^ These off-target effects associated with fluorescence imaging need to be carefully considered by researchers when designing experiments. Bioluminescent indicators such as BLING can offer an orthogonal means to confirm and complement results from fluorescence-based activity studies.

Most importantly, we anticipate BLING will perform well for a variety of molecular imaging applications in deep brain structures and have already successfully used BLING 1.0 to report changes in brain activity while imaging through significantly more material than what can be accomplished with fluorescent techniques. We also believe we can significantly improve on these *in vivo* results when using more sophisticated imaging hardware such as bioluminescent mini scopes that can be used for recording activity in freely behaving animals^35^. This class of sensors will immediately benefit ongoing research efforts to study the mechanisms that give rise to a wide array of neuronal and psychiatric disorders and provide researchers with significantly improved approaches to study neuronal activity at the level of the cell, network, and behaving animals longitudinally. Furthermore, since these reporters produce their own light, they can potentially be used as activators for light-sensitive proteins to carry out a variety of downstream functions within a cell. For example, sensors such as BLING can replace the intact luciferases in current BioLuminescent-OptoGenetic (BL-OG) constructs^20,36^ so the light sensitive ion channels open in response to glutamate resulting in activity dependent excitation or inhibition. Additionally, now that we have created BLING variants with diverse properties such as varying levels of background luminescence and responses to their neurotransmitter (Fig. 6) we can create a variety of neuromodulators based on these sensors. These new BLINGs also present ample opportunity as multiple starting points for further evolution to create new BLINGs with greater responses to glutamate and lower background. These can then be used to improve on prior and ongoing work in neurodegenerative disorders such as spinal cord injury and Parkinson’s Disease by allowing the non-invasive current stimulation of neurons to be dependent on endogenous activity^37,38^.

In conclusion, we were able to successfully engineer a bioluminescent indicator for the neurotransmitter glutamate by adapting sensing domains and split luciferases that have previously been used with success for fluorescent glutamate sensors and bioluminescent calcium sensors. The most optimized BL glutamate sensors we report are already capable of reporting changes in extracellular glutamate and will serve as excellent starting points for engineering derivatives with even higher brightness, dynamic range and for other neurotransmitters. We expect that this indicator will be useful for imaging brain activity within deep brain regions and that we can use this approach to engineer a variety of other neurotransmitter sensors. We also expect this line of neurotransmitter sensors to be adaptable for a variety of highly selective optogenetic actuators that are dependent on a specific neurotransmitter.

## Methods

Sensor Design: The initial three BLING variants were constructed using Glt1, IgK leader and PDFGRβ sequences described in^18^ (GenBank EU42295) synthesized as a Gblock or oligos used for PCR and assembled into a pcDNA 3.1 vector using Gibson Assembly (Neb HiFi) with their respective luciferase fragments. BLING 0.1 consisting of the sbGluc using the split site 105-106^21^, including the 17 AA native secretion signal. BLING 0.2 consisting of NanoLuc split at 66-67^22^, with an Igk leader sequence for cell surface display. BLING 0.3 was created with NanoLuc large and small bits, split at 159-160.^23,24^ (Fig. 2A, Sup. Files 2-4). The additional BLING variants were cloned from synthetic fragments ecoding the 11s variant of NanoLuc and the three affinity variants of the C terminal peptide: peptide 86, peptide 114 and native peptide described in ^24^. HEK cells plated on Poly-D-Lysine coated white 96 well plates, grown to 50-70% confluency were transfected with 0.5 μL Lipofectamine 2000 per mL Opti MEM with 100ng DNA per well using 20μL of the transfection mix per well. Initial testing was done using 5 μM hCTZ (Nanolight Technologies #301) in FluoroBrite media (Thermo), media changed 15 minutes prior to reading to allow the reaction to stabilize and plates read using a BioTek Cytation 5, injecting 10μL of 20 mM glutamate stock into 190 μL media for a final concentration of 1mM.

Library construction and screening: Assembly products were digested with DpnI to eliminate any plasmid template material, eliminating background colonies and electroporated into Top10 cells (Thermo). Colonies were then grown in deep 96 well plates in 1.5mL LB media overnight and miniprepped in 96 well format (Biobasic #B814152-0005). Each plasmid was transfected into HEK cells grown in Poly-D-Lysine coated white 384 well plates in quadruplicate to generate an average for each variant tested. A transfection master mix was prepared with 21mL Opti MEM with 100uL lipofectamine 2000, distributed into 4 96 well PCR plates, 50μL per well and 5 μL DNA from the 96 well mini preps was added to the transfection master mix with mini prep DNA yields ranging from 100-200ng/uL. Testing was done two days later using 5 μM hCTZ in FluoroBrite media, changed 15 minutes prior to reading and plates read using a BioTek Cytation 5, injecting 5μL of 20 mM glutamate stock into 95μL media for a final concentration of 1mM.

Characterization: The top BLING variant from the linker library was termed BLING 1.0 (Addgene plasmid: 171647) and further characterized and compared to the parent construct, BLING 0.2. HEK cells plated on white 96 well plates, grown to 50-70% confluency were transfected with 0.5 μL Lipofectamine 2000 per mL Opti MEM with 100ng DNA per well using 20μL of the transfection mix per well. Measurements were taken with 5 μM hCTZ in FluoroBrite media, changed 15 minutes prior to reading on a Tecan spark for bioluminescence, GCaMP6 and iGluSnFr were taken with a Biotek Cytation 5 using 1 mM glutamate or 5 μM ionomycin respectively. The concentration dependent experiment was done in HBSS with magnesium and calcium with 10 mM HEPES buffer and 1 μM hCTZ as we found these conditions to provide less variability in measurements. Statistical analysis was done using a one-way ANOVA or two way repeated measures with Bonferroni post-hoc n=2-3 per group.

For microscopy experiments, HEK cells seeded in 12 well plates at 8×10^5 per well, transfected the following day using 4μL lipofectamine 2000 in 100μL Opti MEM and 2μg DNA in 100mL Opti MEM, incubated overnight, trypsinized (TrypLE, Thermo) and plated on Poly-D-Lysine coated 18 mm cover slips (NeuVitro), and imaged one to three days later. Imaging was done using a Zeiss A1 Axioscope, 5× 0.17NA objective, Andor iXon 888 EMCCD camera, EM gain of 600, 4×4 binning with an open optical path and microscope within a dark box. Imaging was done in a perfusion chamber with artificial cerebral spinal fluid (ACSF) as described in^36^ using 1 μM furimazine (Promega), heated to 37°C. ACSF was continually perfused followed by ACSF with furimazine followed by ACSF with furimazine and the respective concentration of glutamate followed by a washout with ACSF. The same ROI was used for all concentrations, statistical analysis done using a repeated measures 2-way ANOVA with Bonferroni post-hoc, n=8 per group. Images were analyzed using ImageJ for background subtraction and despeckling to reduce noise, ROIs selected manually for quantification.

All animal work was approved by Michigan State Universities Institutional Animal Care and Use Committee. BLING 1.0 was cloned into an AAV expression vector with a synapsin promoter for expression in neurons and AAV was made and purified in house as previously described^38^. Sprague Dawley rats were injected with 2μL of 0.5×1012 copies/mL at rate of 0.5μL per minute with a 33G world precision syringe at 0mm AP 3.5mm R, and 2mm ventral of the cortical surface and the needle was left in place for 5 minutes after infusion and slowly retracted. Imaging was done at least three weeks later with an IVIS spectrum (Perkin Elmer). Animals were injected with water soluble *in vivo* h-CTZ (Nanolight Technologies) 2mg/kg intraperitoneally one to two hours prior to imaging and checked for the onset of bioluminescence. Once bioluminescence was detectable, the incision was reopened and a custom made cannula was inserted through the small burr hole from the prior surgery for bicuculline administration. Then a baseline recording of ten minutes was acquired and 10μL of 10mM bicuculline was slowly injected over five minutes. Images were captured using large binning, fstop of one with one minute exposures. Images were analyzed with Living Image software with an ROI over the entire skull.

## Acknowledgments

Figures contain material created with BioRender. A.A.G acknowledges financial support from the NSF 2027113, NIH/NINDS: R01-NS098231; R01-NS104306 NIH/NIBIB: R01 EB031008; R01EB030565; R01EB031936 and P41-EB024495. E.D.P was supported Perkin Elmer Postdoctoral Fellowship.

## References

1 Lu, R. et al. Rapid mesoscale volumetric imaging of neural activity with synaptic resolution. Nature Methods, doi:10.1038/s41592-020-0760-9 (2020).

2 Xu, Y., Zou, P. & Cohen, A. E. Voltage imaging with genetically encoded indicators. Curr Opin Chem Biol 39, 1–10, doi:10.1016/j.cbpa.2017.04.005 (2017).

3 Chamberland, S. et al. Fast two-photon imaging of subcellular voltage dynamics in neuronal tissue with genetically encoded indicators. Elife 6, doi:10.7554/eLife.25690 (2017).

4 St-Pierre, F. et al. High-fidelity optical reporting of neuronal electrical activity with an ultrafast fluorescent voltage sensor. Nat Neurosci 17, 884–889, doi:10.1038/nn.3709 (2014).

5 Tian, L. et al. Imaging neural activity in worms, flies and mice with improved GCaMP calcium indicators. Nat Methods 6, 875–881, doi:10.1038/nmeth.1398 (2009).

6 Patriarchi, T. et al. Ultrafast neuronal imaging of dopamine dynamics with designed genetically encoded sensors. Science 360, doi:10.1126/science.aat4422 (2018).

7 Marvin, J. S. et al. An optimized fluorescent probe for visualizing glutamate neurotransmission. Nat Methods 10, 162–170, doi:10.1038/nmeth.2333 (2013).

8 Raghuram, A. et al. Determining the Depth Limit of Bioluminescent Sources in Scattering Media. bioRxiv, 2020.2004.2021.044982, doi:10.1101/2020.04.21.044982 (2020).

9 Wilson, T. & Hastings, J. W. BIOLUMINESCENCE. Annual Review of Cell and Developmental Biology 14, 197–230, doi:10.1146/annurev.cellbio.14.1.197 (1998).

10 Martini, S. & Haddock, S. H. D. Quantification of bioluminescence from the surface to the deep sea demonstrates its predominance as an ecological trait. Scientific Reports 7, 45750, doi:10.1038/srep45750 (2017).

11 Bonora, M. et al. Subcellular calcium measurements in mammalian cells using jellyfish photoprotein aequorin-based probes. Nat Protoc 8, 2105–2118, doi:10.1038/nprot.2013.127 (2013).

12 Granatiero, V., Patron, M., Tosatto, A., Merli, G. & Rizzuto, R. Using targeted variants of aequorin to measure Ca2+ levels in intracellular organelles. Cold Spring Harb Protoc 2014, 86–93, doi:10.1101/pdb.prot072843 (2014).

13 Thomou, T. et al. Adipose-derived circulating miRNAs regulate gene expression in other tissues. Nature 542, 450–455, doi:10.1038/nature21365 (2017).

14 Iwano, S. et al. Single-cell bioluminescence imaging of deep tissue in freely moving animals. Science (New York, N.Y.) 359, 935–939, doi:10.1126/science.aaq1067 (2018).

15 Abe, M. et al. Near-Infrared Bioluminescence Imaging with a through-Bond Energy Transfer Cassette. Chembiochem: A European Journal of Chemical Biology, doi:10.1002/cbic.201900149 (2019).

16 Patriarchi, T. et al. Ultrafast neuronal imaging of dopamine dynamics with designed genetically encoded sensors. Science, eaat4422, doi:10.1126/science.aat4422 (2018).

17 Sun, F. et al. A Genetically Encoded Fluorescent Sensor Enables Rapid and Specific Detection of Dopamine in Flies, Fish, and Mice. Cell 174, 481–496.e419, doi:10.1016/j.cell.2018.06.042 (2018).

18 Hires, S. A., Zhu, Y. & Tsien, R. Y. Optical measurement of synaptic glutamate spillover and reuptake by linker optimized glutamate-sensitive fluorescent reporters. Proceedings of the National Academy of Sciences 105, 4411, doi:10.1073/pnas.0712008105 (2008).

19 Welsh, J. P., Patel, K. G., Manthiram, K. & Swartz, J. R. Multiply mutated Gaussia luciferases provide prolonged and intense bioluminescence. Biochemical and Biophysical Research Communications 389, 563–568, doi:https://doi.org/10.1016/j.bbrc.2009.09.006 (2009).

20 Park, S. Y. et al. Novel luciferase-opsin combinations for improved luminopsins. J Neurosci Res 98, 410–421, doi:10.1002/jnr.24152 (2020).

21 Kim, S. B., Sato, M. & Tao, H. Circularly Permutated Bioluminescent Probes for Illuminating Ligand-Activated Protein Dynamics. Bioconjugate Chemistry 19, 2480–2486, doi:10.1021/bc800378a (2008).

22 Suzuki, K. et al. Five colour variants of bright luminescent protein for real-time multicolour bioimaging. Nat Commun 7, 13718, doi:10.1038/ncomms13718 (2016).

23 Farhana, I., Hossain, M. N., Suzuki, K., Matsuda, T. & Nagai, T. Genetically Encoded Fluorescence/Bioluminescence Bimodal Indicators for Ca2+ Imaging. ACS Sensors 4, 1825–1834, doi:10.1021/acssensors.9b00531 (2019).

24 Dixon, A. S. et al. NanoLuc Complementation Reporter Optimized for Accurate Measurement of Protein Interactions in Cells. ACS Chemical Biology 11, 400–408, doi:10.1021/acschembio.5b00753 (2016).

25 Cheng, K. P., Kiernan, E. A., Eliceiri, K. W., Williams, J. C. & Watters, J. J. Blue Light Modulates Murine Microglial Gene Expression in the Absence of Optogenetic Protein Expression. Scientific Reports 6, 21172, doi:10.1038/srep21172 (2016).

26 Picot, A.et al. Temperature Rise under Two-Photon Optogenetic Brain Stimulation. Cell Reports 24, 1243–1253.e1245, doi:10.1016/j.celrep.2018.06.119 (2018).

27 Podgorski, K. & Ranganathan, G. Brain heating induced by near-infrared lasers during multiphoton microscopy. Journal of Neurophysiology 116, 1012–1023, doi:10.1152/jn.00275.2016 (2016).

28 Stockley, J. H. et al. Surpassing light-induced cell damage in vitro with novel cell culture media. Scientific Reports 7, 849, doi:10.1038/s41598-017-00829-x (2017).

29 Owen, S. F., Liu, M. H. & Kreitzer, A. C. Thermal constraints on in vivo optogenetic manipulations. Nature Neuroscience 22, 1061–1065, doi:10.1038/s41593-019-0422-3 (2019).

30 Inagaki, S. et al. Genetically encoded bioluminescent voltage indicator for multi-purpose use in wide range of bioimaging. Scientific Reports 7, 42398, doi:10.1038/srep42398 (2017).

31 Jin, L. et al. Single action potentials and subthreshold electrical events imaged in neurons with a novel fluorescent protein voltage probe. Neuron 75, 779–785, doi:10.1016/j.neuron.2012.06.040 (2012).

32 Xu, Y., Zou, P. & Cohen, A. E. Voltage imaging with genetically encoded indicators. Current Opinion in Chemical Biology 39, 1–10, doi:10.1016/j.cbpa.2017.04.005 (2017).

33 Hochbaum, D. R. et al. All-optical electrophysiology in mammalian neurons using engineered microbial rhodopsins. Nature methods 11, 825–833, doi:10.1038/nmeth.3000 (2014).

34 Abdelfattah, A. S. et al. A Bright and Fast Red Fluorescent Protein Voltage Indicator That Reports Neuronal Activity in Organotypic Brain Slices. Journal of Neuroscience 36, 2458–2472, doi:10.1523/JNEUROSCI.3484-15.2016 (2016).

35 Celinskis, D. et al. Miniaturized Devices for Bioluminescence Imaging in Freely Behaving Animals. Annu Int Conf IEEE Eng Med Biol Soc 2020, 4385–4389, doi:10.1109/embc44109.2020.9175375 (2020).

36 Berglund, K. et al. Luminopsins integrate opto- and chemogenetics by using physical and biological light sources for opsin activation. Proceedings of the National Academy of Sciences 113, E358, doi:10.1073/pnas.1510899113 (2016).

37 Zenchak, J. R. et al. Bioluminescence-driven optogenetic activation of transplanted neural precursor cells improves motor deficits in a Parkinson’s disease mouse model. J Neurosci Res 98, 458–468, doi:10.1002/jnr.24237 (2020).

38 Petersen, E. D. et al. Restoring Function After Severe Spinal Cord Injury Through Bioluminescence-Driven Optogenetics. bioRxiv, 710194, doi:10.1101/710194 (2019).

