## Supplemental material for "Bioluminescent Genetically Encoded Glutamate Indicator for Molecular Imaging of Neuronal Activity"

Supplementary – Petersen

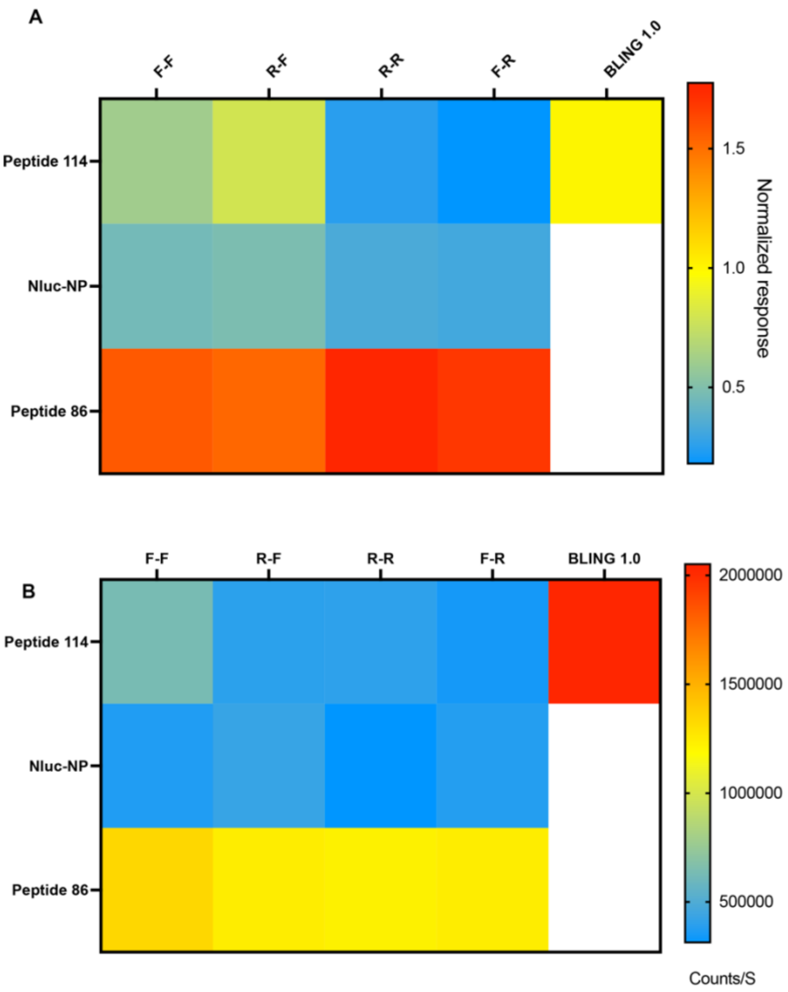


Figure S1. A. Response of NanoBiT based BLING variants consisting of three different C terminal variants of different affinities to the N terminal luciferase in combination with either flexible or rigid linkers normalized to BLING 1.0. B. average brightness of these twelve BLING variants compared to BLING 1.0.
